# Sociality Drives Sex Differences in Auditory Brainstem Processing Across Rodent Species

**DOI:** 10.64898/2026.06.09.731227

**Authors:** Luberson Joseph, Desi M. Joseph, Naleyshka Colon-Rivera, Fabio A. Machado, Elizabeth A. McCullagh

**Author notes:** Corresponding author’s.

## Abstract

Although social behavior has been predicted to correlate with increased complexity in vocal signals, its relationship with auditory sensitivity between the sexes remains poorly understood. Here, we used phylogenetic comparative analyses to examine sex-specific differences in auditory processing across rodent species representing different social lifestyle strategies. We detected significant sex differences in click and frequency evoked auditory brainstem response (ABR) thresholds across sociality, with social female rodents exhibiting lower thresholds than solitary males. Males generally exhibited higher ABR wave I and IV amplitude ratios than females, whereas interpeak latencies were similar between the sexes across social groups. Females exhibited significantly higher binaural interaction component (BIC) relative amplitudes and faster BIC normalized latencies than males across tested interaural time differences (ITDs). Together, these findings demonstrate that sociality plays an important role in shaping differences in auditory physiology between male and female rodents and highlights the potential influence of social behavior on the evolution of mammalian auditory systems.

## INTRODUCTION

Social organization (hereafter sociality) is diverse in form and function and is widespread among small mammals including solitary-living, pair-living or monogamous, group-living, and eusocial-living^1,2^. These distinct forms of sociality foster different communication strategies that enable individuals to signal conspecifics for building and maintaining steady social structures, coordinating activities, avoiding predators, and securing mates^3–7^. Within and across social systems, small mammals, like members of the order Rodentia, rely on a wide range of sensory modalities to communicate, including olfactory, tactile, visual, and auditory cues^3–6^. Among these modalities, auditory communication is uniquely suited for precise, rapid, and long-distance information sharing and also plays an essential role in mediating social interactions^7^, making it a powerful system for examining how sociality may shape the evolution and neural processing of signals.

In recent years, comparative studies have increasingly sought to elucidate the relationship between sociality and acoustic communication^8–10^. Indeed, several hypotheses have been proposed to explain the coevolution of sociality and vocal communication across tetrapoda^10,11^. The Social Complexity Hypothesis for Communication (SCHC) predicts that species living in more complex social environments exhibit a higher diversity of vocal signals phenotypes than solitary-living species^11^, and it therefore stands to reason that natural and sexual selection should favor corresponding tuning of receiver auditory sensitivity to socially relevant signals^12,13^. This hypothesis has been tested in various mammalian taxa with diverse social systems (e.g., giant otter, whales, meerkats, non-human primates, bats, slender mongooses, and rodents), and compelling evidence suggests that increases in social complexity are associated with greater vocal repertoire size and structural signal complexity^14–20^. In African striped mice (*Rhabdomys pumilio*) for instance, larger social groups are associated with enhanced vocal diversity, with individuals vocalizing more frequently in the presence of familiar conspecifics and altering their vocal repertoires when communicating in the presence of unfamiliar individuals^20^. However, most prior studies have been exclusively centered on the signaler side of acoustic communication, while largely neglecting the receiver side. Accordingly, this disparity limits our ability to fully evaluate communication as an integrated signaler-receiver system and obscures the role of auditory processing constraints and adaptation in the evolution of social communication. Comparative studies are therefore critical for illuminating the connection between sociality and acoustic signal processing across vertebrates differing in social lifestyle strategies.

Although the SCHC has primarily been applied to vocal signal production^14–19^, its framework can be extended to the receiver’s side by relating auditory processing ability to social lifestyle organizations. Rodents represent an ideal comparative model system for such analyses owing to their exceptional diversity in social structures, mating systems, and parental care strategies. Previous studies in primates and rodents indicate that social species exhibit enhanced auditory sensitivity and expanded high-frequency hearing relative to solitary living species^10,21^, suggesting that social environments can shape receiver traits. At the same time, substantial sex differences in auditory processing have been documented across rodents, with females generally exhibiting more sensitive hearing and producing higher frequency vocalizations than males^22–25^. For instance, studies in several mouse strains (CBA/J, CBA/CaJ, etc.) have demonstrated that female mice exhibit lower frequency evoked thresholds than males^22,26^. Similarly, female rats (Brattleboro, long-Evans, etc.,) exhibit greater auditory sensitivity across multiple frequencies compared to males^27^. Sex differences in auditory physiology have also been reported in the solitary hispid pocket mouse (*Chaetodipus hispidus*) and socially monogamous prairie voles (*Microtus ochrogaster*), where females of both species generally exhibit lower click and frequency evoked thresholds^24,28,29^, increased amplitude of peripheral ABR waveforms I and II, and longer latency of waveforms III and IV compared to males^29^. These findings suggest that auditory processing can differ between the sexes in multiple ways, potentially reflecting differences in parental investment, communication roles, social, and reproductive behavior.

However, it remains unclear whether and how these male-female differences in auditory physiology vary across species that differ in social lifestyle strategies. Accordingly, comparing auditory processing ability between male and female rodents with diverse social structures and parental care strategies provides a powerful opportunity to determine whether the magnitude and pattern of sex differences in auditory processing vary with social lifestyle strategies. Here, we used auditory brainstem response (ABR) recordings and phylogenetically controlled analyses to examine whether sociality drives differences in auditory processing abilities between the sexes in nine wild-caught rodent species. We compared solitary and social rodents using click and frequency evoked thresholds, amplitude ratios, interpeak latencies, binaural interaction component (BIC) relative amplitudes, and BIC normalized latencies. We hypothesized that sex differences in auditory processing would vary with sociality because social communication and parental care impose distinct sensory demands on males and females. Specifically, we predicted that social species would exhibit greater auditory sensitivity than solitary species, reflected by lower click and frequency evoked ABR thresholds, particularly within the 8-32 kHz range of peak rodent hearing sensitivity. We further predicted that sex differences in temporal and binaural processing would be more pronounced in social species than solitary species, with social species exhibiting larger amplitude ratios, shorter interpeak latencies, higher BIC relative amplitudes, and faster BIC normalized latencies compared to solitary species.

## MATERIALS AND METHODS

### Field Sampling and Ethical Notes

A total of 180 rodents representing 9 species (10 males and 10 females per species, Table 1) were included in this study. These individuals consist of 90 males previously reported in Joseph and colleagues^21^ and 90 newly collected females used here for sex-based comparative analyses. Individuals were live trapped using aluminum Sherman non-folding traps (7.62 cm x 7.62 cm x 25.4 cm; H.B Sherman Traps, Inc., Tallahassee, FL) baited with whole grain oats and peanut butter. Trapping was conducted between June 2022 and July 2025 across six locations in Oklahoma (Selman living laboratory, Stillwater, Tulsa, James Collin, Sandy Sanders and Packsaddle wildlife management areas) and one location in Kansas (Kansas University field station, Lawrence, Fig 1). Rodents were collected along two trap-lines consisting of 10 traps each, spaced 5-10 m apart. To minimize thermal stress, traps were positioned beneath vegetation or brush from mid-May to end of November (warm months), whereas 2-3 cotton balls were added for warmth from December to end of April (cold months) of each year for insulation. Traps were deployed at dusk (18:00 – 20:00 h) and inspected the following morning at sunrise (6:00 – 9:00 h). Upon capture, individuals were temporarily placed in unsealed plastic bags for visual identification of species and sex according to Caire and colleagues^30^. Age was estimated for each individual using body mass criteria derived from the primary literature, and animals were categorized as juveniles, subadults, or adults (Table 1). Animals were subsequently housed individually in disposable plastic mouse cages (Innovive: 37.34 x 23.37 x 13.97 cm) and transported to the laboratory for ABR recordings. All animal collection and transportation adhered to the ethical guidelines of the American Society of Mammalogists (ASM)^31^, the Oklahoma Department of Wildlife Conservation (ODWC), the Kansas Department of Wildlife and Parks (KDWP), the Selman Living Laboratory (SLL) of the University of Central Oklahoma (UCO), the Kansas University (KU) field station, and Oklahoma State University (OSU). All experimental protocols were approved by the Institutional Animal Care and Use Committee (IACUC) at OSU in Stillwater, Oklahoma (protocol number: 22-09).

**Table 1:** Scientific name, social structure, age group, body mass range (g), sample size by age class (juvenile, subadult, and adult), and corresponding references for each species included in the study.

| Scientific name | Social structure | Age group | Body mass range (g) | Male sample size (Juvenile, subadult, adult) | Female sample size (Juvenile, subadult, adult) | References |
| --- | --- | --- | --- | --- | --- | --- |
| *Mus musculus* | Social | Juvenile<br>Subadult<br>Adult | < 10<br>10-14<br>> 14 | 10 (2, 3, 5) | 10 (0, 4, 6) | 38-40 |
| *Rattus norvegicus* | Social | Juvenile<br>Subadult<br>Adult | < 100<br>100-120<br>> 120 | 10 (3, 1, 6) | 10 (3, 0, 7) | 37,41,42 |
| *Microtus ochrogaster* | Social | Juvenile<br>Subadult<br>Adult | < 20<br>20-30<br>> 30 | 10 (1, 2, 7) | 10 (0, 2, 8) | 43-45 |
| *Onychomys leucogaster* | Social | Juvenile<br>Subadult<br>Adult | < 17<br>17-25<br>> 25 | 10 (0, 1, 9) | 10 (0, 1, 9) | 36,46,47 |
| *Sigmodon hispidus* | Solitary | Juvenile<br>Subadult<br>Adult | < 80<br>80-110<br>> 110 | 10 (3, 3, 4) | 10 (2, 4, 4) | 48-50 |
| *Chaetodipus hispidus* | Solitary | Juvenile<br>Subadult<br>Adult | < 30<br>30-35<br>> 35 | 10 (6, 0, 4) | 10 (6, 1, 3) | 34,51,52 |
| *Peromyscus leucopus* | Solitary | Juvenile<br>Subadult<br>Adult | < 13<br>13-18<br>> 18 | 10 (1, 3, 6) | 10 (0, 6, 4) | 35,53-55 |
| *Dipodomys ordii* | Solitary | Juvenile<br>Subadult<br>Adult | < 40<br>40-52<br>> 52 | 10 (0, 3, 7) | 10 (2, 3, 5) | 33,56,57 |
| *Neotoma floridana* | Solitary | Juvenile<br>Subadult<br>Adult | < 100<br>100-140<br>> 140 | 10 (1, 2, 7) | 10 (1, 1, 8) | 58,59 |

**Fig 1:**
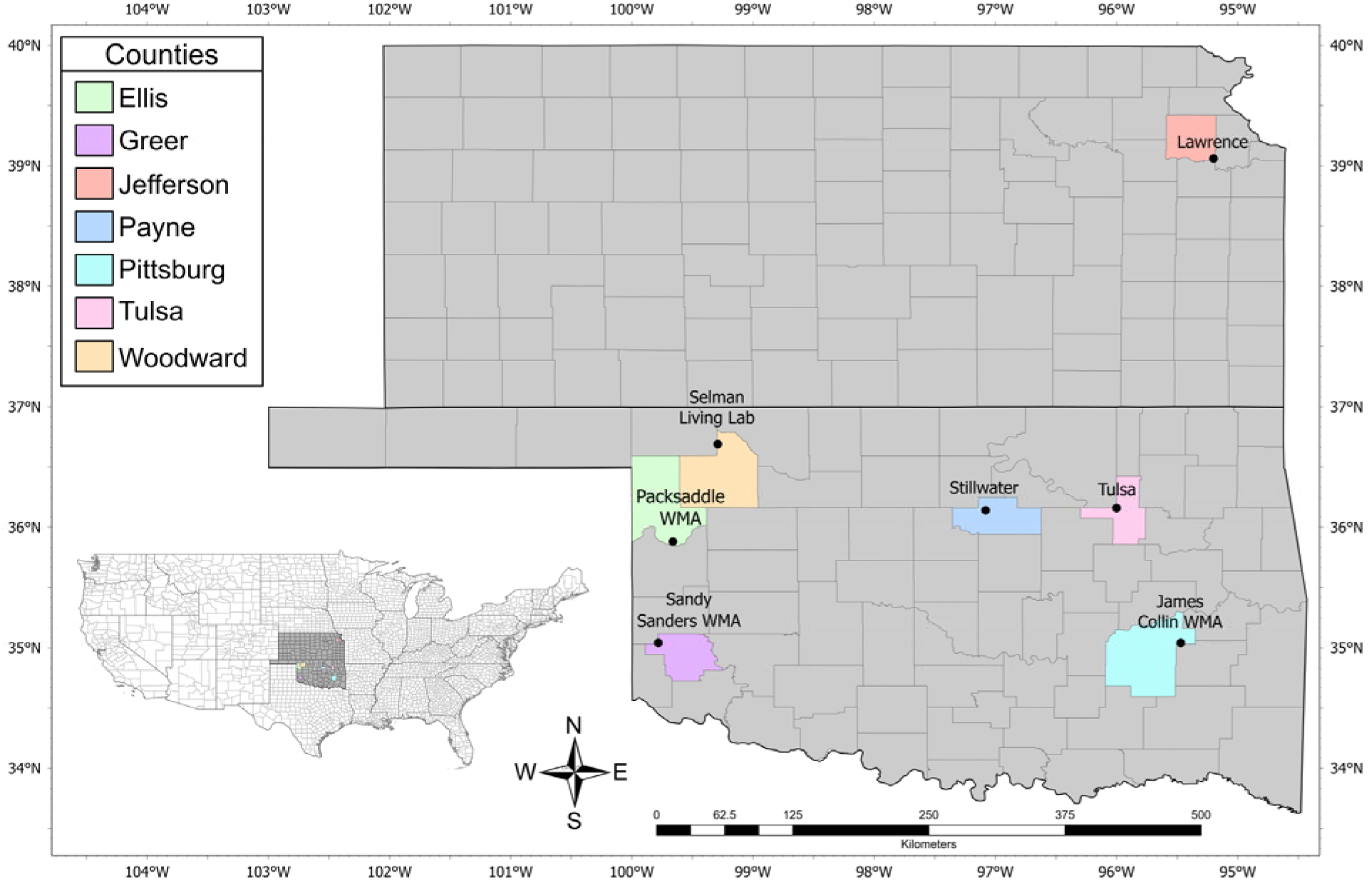
Map showing the seven sampling locations in the states of Oklahoma and Kansas. Each sampling location is indicated by a black circle, and black lines denote state and county boundaries. Colored counties represent areas where sampling occurred in both Oklahoma and Kansas.

### Sociality Classification

Species were classified into two social categories (i.e., solitary and social) based on patterns of parental care and breeding structure reported in the primary literature^32^. Solitary species were characterized by exclusively maternal care and spatially separated breeding females, with social interaction largely restricted to mating and mother-offspring relationships^33–35^. Social species included both monogamous and group-living rodents. We combined these two social systems into a single category because our previous work found no overall differences in hearing thresholds or ABR wave metrics between them in male individuals^21^. Social species were characterized by stable social associations, including long-term pair bonds or group-living with shared home ranges or breeding sites, often accompanied by cooperative interactions, mutual tolerance, and extensive spatial overlap^35–37^. Species were assigned to specific social category based on consistent descriptions of social organization from field-based studies and reflect the predominant social system reported for each species rather than population-specific variation across social contexts^33–37^.

### ABR Recordings

We recorded ABRs to examine sex differences in auditory processing across species with differing social lifestyle strategies (Fig 2A) using procedures as previously described^24,29,60–62^. Animals were intraperitoneally anesthetized to an areflexive state with a mixture of ketamine (60 mg/kg) and xylazine (10 mg/kg) and kept sedated with ketamine (25 mg/kg) and xylazine (12 mg/kg) dose supplements throughout the experiment. To monitor the anesthetic state of the tested individual, we confirmed toe pinch reflex responses and breathing rates every six minutes for the duration of the recording. ABR measurements were recorded inside a sound-attenuating chamber (Noise Barriers, Lake Forest, IL, USA) lined with SONEX acoustic foam in the interior (Illbruck Inc., Minneapolis, MN). Prior to testing, animals were placed on a heating pad to maintain a body temperature of 37°C using a water pump heating pad (25.5 cm x 17.8 cm; HTP-1500 Heat Therapy Pump; Kent Scientific, Torrington, CT). Three subdermal needle electrodes (Viasys Healthcare, Madisson, Wi, USA) were placed under the skin between the pinnae at the vertex of the head (active electrode), behind the apex of the nape (reference electrode), and the right back leg (ground electrode) to measure and visualize the ABR signals. Tucker-Davis Technologies (TDT, Alachua, FL, USA) RA4LI head stage, a multi I/O processor RZ5, and an RA16PA preamplifier attached to an Acer PC running custom Python software positioned outside of the ABR chamber were used to coordinate stimuli presentations and visualize evoked potential response acquisition.

**Fig 2A:**
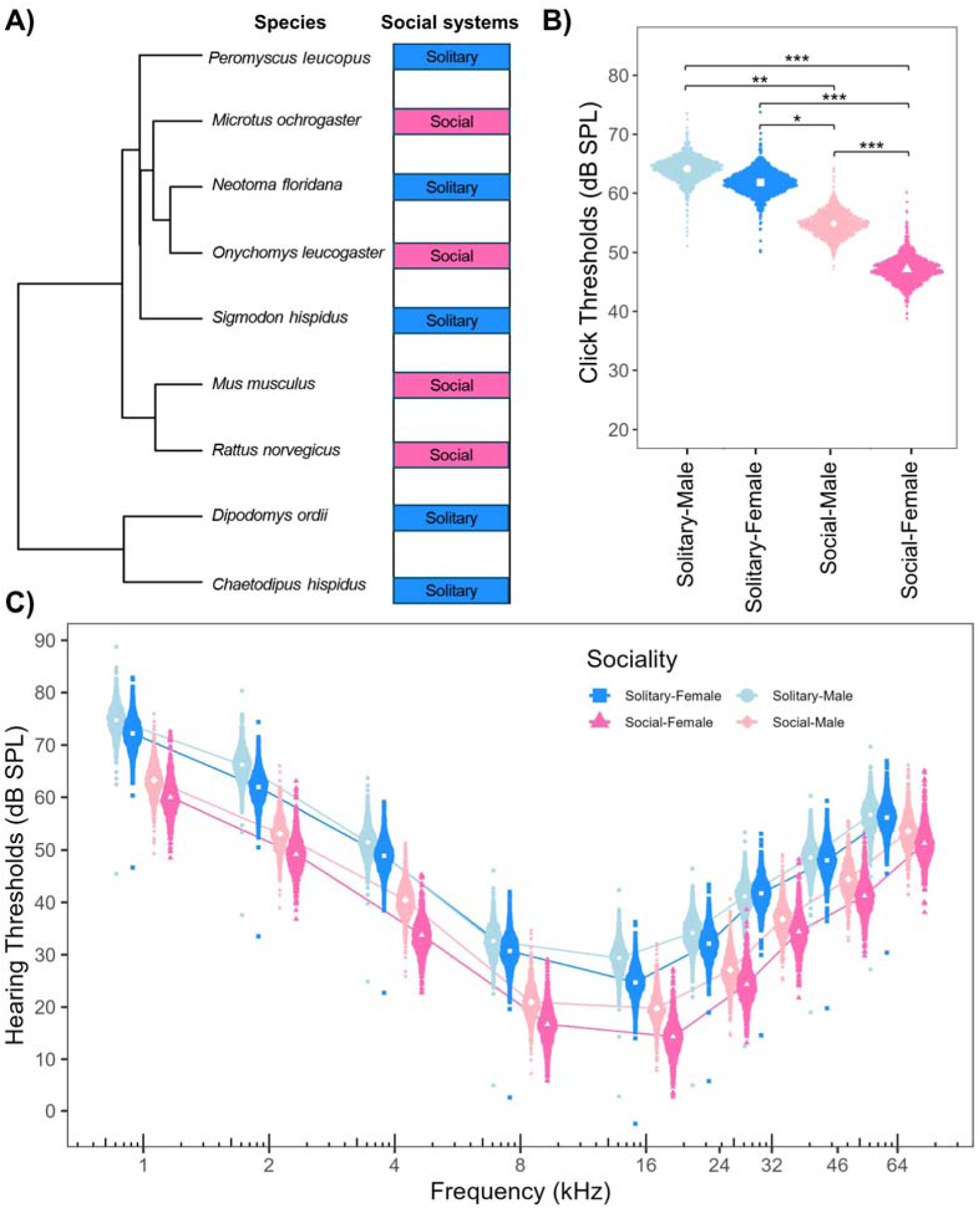
Phylogeny used for the analysis of wild-caught rodent’s ABR data. The phylogenetic tree was reconstructed from Upham and colleagues using basic local alignment search tools from the National Center for Biotechnology information nucleotide database^70^. Colors represent different social systems across species: blue = solitary, pink = social. **B** represents mean click thresholds between male and female wild-caught rodents across social categories. **C** shows mean auditory brainstem response thresholds between the sexes across tested frequencies. Significant sex differences were detected in frequency evoked and click response thresholds across social groups. Lines (blue, and pink) connect mean responses at each frequency to illustrate possible trends across frequencies. [(n = 180, 20 individuals per species (10 males and 10 females)]. Asterisks represent Bayesian statistical differences across social groups with P < 0.05 *, P ≤ 0.01 **, P ≤ 0.001 ***.

### Acoustic Stimuli

Acoustic stimuli were delivered either through Tucker-Davis Technology multi-field magnetic speakers (TDT MF-1) for clicks and frequencies ranging from 1 to 24 kHz or TDT coupled electrostatic speakers (TDT EC-1) for frequencies between 32 and 64 kHz. Speakers were positioned at the animal’s left and right ear canals for close-field recording. Speakers were coupled with custom ear bars fitted with Etymotic ER-7C probe microphones (Etymotic Research Inc., Elk Grove Village, IL) that we used for in-ear calibration prior to recording. Stimuli consisted of clicks (0.1 ms) and tones [4 ms total with 1 ms rise and 1 ms fall ramps (2 ms ± 1 ms rise/fall ramps)]. ABR recording data were filtered using a second-order bandpass filter (50-3000 Hz) and averaged over a 10-12 ms window across 500 – 1000 stimulus repetitions. Sound stimuli were presented to the anesthetized animal with an interstimulus interval of 30 ms and a standard deviation of 5 ms at different intensity (90, 80, 70, 60, 50, 40, 30, 20, 10, 5 dB SPL), starting with the largest sound levels. Stimuli generation was performed at a 97656.25 Hz sampling rate using a TDT RP2.1 real-time processor controlled via custom Python software^63^.

### ABR Thresholds

Two methods were used to estimate auditory thresholds of rodents between the sexes. Click evoked threshold was first assessed by gradually reducing the intensity of the click stimulus in 5-10 dB SPL steps until the ABR waveforms were no longer visible. Click threshold was denoted as the difference in intensity between the last level where ABR waveforms were detected and the level at which they disappeared (i.e., if ABR waveforms were visible at 50 dB SPL and no longer detectable at 40 dB SPL, the click threshold was estimated to be 45 dB SPL for this animal). We next determined frequency evoked thresholds with pure tone stimuli to analyze the audiogram across frequencies (1, 2, 4, 8, 16, 24, 32, 46, and 64 kHz) and intensities [10 - 90 dB sound pressure level (SPL)] delivered to the anesthetized animal. ABR frequency thresholds were measured as the intermediate sound level between the lowest level of stimulation that produced a response different from noise and the level through which no responses were visible by both left and right monaural stimuli^64–66^.

### ABR Amplitude Ratio and Interpeak Latency

We independently presented broadband click stimuli to the right and left pinnae of the anesthetized rodent to elicit monaural evoked potentials across tested intensities (50 - 90 dB SPL). Absolute latency (i.e., the time between the onset of the sound stimulus and the point of maximum amplitude of each corresponding wave) and absolute amplitude (i.e., the voltage from peak to trough difference) were measured for wave I and wave IV from the ABR data acquired from the left and right pinnae across tested intensities. We further estimated amplitude ratio and interpeak latency (relative measures accounting for inter-individual variability) for each species to make sex-specific comparisons between social systems. Amplitude ratio was determined as the ratio of wave IV amplitude to wave I amplitude for the left and right pinnae at each intensity (50 – 90 dB SPL), whereas inter-peak latency was calculated as the difference between the peak latencies of wave I and IV at each intensity level for both pinnae. We then used the average of the calculated amplitude ratio and inter-peak latency data at each intensity for comparisons between the sexes between solitary and social rodents^24,60,67^.

### Binaural ABR

Broadband alternating polarity click stimuli were presented to both pinnae of the sedated animal simultaneously or with an ITD (ranging from 0.0 to 2.0 ms in 0.5 steps) at 90 dB SPL to elicit evoked binaural auditory responses. To assess binaural auditory sensitivity between the sexes, the BIC was quantified and adjusted to the zero baseline of each recording using the RMS of the waveform to ensure similar alignment across measurements. The BIC was calculated by taking the sum of the monaural left and right absolute latency and absolute amplitude responses and subtracting from the binaural evoked ABR recordings^63,68^. BIC was characterized as the prominent negative peak (DN1) occurring around wave IV of the binaural ABR following the subtraction of the monaural and binaural ABR responses^29^. We next calculated BIC latency and BIC amplitude using custom Python Software, with latency measured as the time point corresponding to the peak of the DN1 component, while amplitude was measured referenced to the recording baseline as described by New and colleagues^29^. Previous studies have shown that as ITD increases, DN1 normalized latency gets longer, while DN1 relative amplitude decreases^24,60,63,68^. To quantify ITD-dependent changes, DN1 latency and amplitude were expressed relative to their values at 0 ms ITD (BIC normalized latency shift = latency at a given ITD minus latency at 0 ms, BIC relative amplitude = amplitude at a given ITD divided by amplitude at 0 ms x 100)^63^. This normalization accounts for potential size differences between sexes and species within each social group. We further used the average BIC normalized DN1 latency shift and BIC relative amplitude values to explore sex-specific difference in binaural auditory sensitivity as a function of ITD between solitary and social rodents.

### Statistical Analyses

Phylogenetic comparative analyses were conducted to compare hearing thresholds and ABR wave characteristics between the sexes using Bayesian phylogenetic generalized linear mixed models (PGLMMs) to account for shared evolutionary history among species. A time-calibrated species-level phylogeny was obtained by pruning the mammalian phylogeny of Upham and colleagues using the R package APE to incorporate only species sampled in this work^69,70^. We fit individual PGLMM models for each auditory response variable, including frequency evoked thresholds, click thresholds, amplitude ratios, interpeak latencies, BIC relative amplitudes, and BIC normalized latencies. All response variables were modeled assuming Gaussian error distributions. Fixed effects variables including sex (male and female), stimulus intensity (50-90 dB SPL), sociality (solitary and social), frequency (1-64 kHz), ITD (0.0-2.0 ms), and their interaction terms where specify (See supplementary table 1). Across models, ITD, frequency, and intensity were all modelled as categorical variables to correspond to the discrete stimulus conditions specified in the experimental design. Phylogenetic relatedness and individual identity were treated as random effects to account for the repeated measure structure of the ABR data recorded. We did not include age and mating status in any statistical analyses owing to uncertainty in mating status and the inherent variability and indirect measure of age estimation in wild-caught rodents. We quantified phylogenetic signal as phylogenetic heritability (*H*^2^), defined as the proportion of total variance explained by phylogenetic effects relative to residual variance, following Hadfield and Nakagawa^71^ :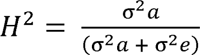, where σ^2^*a* denotes phylogenetic variance and σ^2^*e* denotes residual variance from the best supported model.

Posterior distributions were estimated using Markov Chain Monte Carlo (MCMC) sampling implemented in MCMCglmm^72^. The default diffuse Gaussian prior was applied to fixed effects (mean = 0, variance = 10^10^), and the weakly informative inverse-Gamma prior (V = 1) for random effects with shape and scale parameters set to 0.001. We ran individual models with three independent chains, and convergence was assessed using the Gelman-Rubin test, with a scale reduction factor of one indicated satisfactory convergence and identical chains^73–75^. Separate models were run for 260,000 interactions, with a burn-in of 10,000 and thinning every 250 steps to decrease autocorrelation. To identify the best-fit model for each response variable, we compared four candidate models (null, additive, full model, and a full model with body mass as a covariate variable). The null model included random effects (phylogeny and individual ID) and a single predictor variable (frequency, intensity, or ITD). The additive model included the random effects (phylogeny and individual ID) and three predictors as additive terms (sex + sociality + frequency, sex + sociality + intensity, or sex + sociality + ITD). The full model included the random effects (phylogeny and individual ID) and interaction terms between the predictors (sex * sociality * frequency, sex * sociality * intensity, or sex * sociality * ITD). The full model with body mass (BM) as a covariate was used to account for inter-individual differences in body mass. Model selection was performed by comparing null, additive, full interaction model, and full interaction model and body mass using Devariance Informative Criterion (DIC) to select the best fit model for each response variable (Supplementary table 1). The simplest model was selected in scenario where differences in DIC score across model were small (DIC < 2). Full models were strongly supported for most response variables, except for interpeak latency where the additive model was the best fit. We considered a fixed effect to be significant when the 95% credible intervals did not overlap with zero and when the Bayesian P_MCMC_ estimate was less than 0.05. When there was an overall difference of a tested variable, we used Bayesian contrasts (two-tailed comparisons of posterior distributions) for pairwise comparisons. All statistical analyses were performed in R^76^, and figures were generated using ggplot2^77^ and Python^78^.

## RESULTS

### Click Threshold Sensitivity

We detected significant sex differences in click threshold when data were analyzed across species independent of sociality (PGLMM: Phylogenetic Heritability (H^2^) = 0.47, Posterior Mean (PM) = 5.17, 95% Credible Interval (95% CrI) = 2.43 to 7.76, P_MCMC_ = 0.0003). Male rodents exhibited a mean click threshold of 61.62 dB SPL, whereas females exhibited lower mean thresholds of 56.45 dB SPL (Supplementary Fig 1). When data were analyzed for sex differences within social categories, no significant differences in click evoked thresholds were detected between solitary male and solitary female rodents (PGLMM: H^2^ = 0.05, PM = 2.31, 95% CI = -1.73 to 6.46, P_MCMC_ = 0.258). In contrast, social males exhibited higher click thresholds than social females (PGLMM: H^2^ = 0.05, PM = 7.77, 95% CrI = 3.97 to 11.99, P_MCMC_ < 0.001). When comparing between social groups, PGLMM analyses revealed that solitary males exhibited higher click thresholds than both social males (PGLMM: H^2^ = 0.05, PM = 9.23, 95% CrI = 3.92 to 13.5, P_MCMC_ = 0.006) and social females (PGLMM: H^2^ = 0.05, PM = 17, 95% CrI = 12.17 to 21.3, P_MCMC_ < 0.001). Similarly, solitary females exhibited higher click thresholds than social males (PGLMM: H^2^ = 0.05, PM = 6.92, 95% CrI = 1.48 to 11.62, P_MCMC_ = 0.022) and social females (PGLMM: H^2^ = 0.05, PM = 14.69, 95% CrI = 9.26 to 19.66, P_MCMC_ < 0.001).

### Hearing Sensitivity

We constructed audiograms for each species across a range of sound intensities and frequencies to assess sex differences in hearing sensitivity as previously described^24,60,61^. Across all species, independent of sociality, both sexes exhibited the lowest auditory thresholds between 8 and 24 kHz, indicating peak hearing sensitivity within this frequency range (Fig 2C). PGLMM analyses with Bayesian MCMC sampling revealed significant sex differences in hearing sensitivity across species, with females exhibiting lower auditory evoked thresholds than males when considering all tested frequencies (PGLMM: H^2^ = 0.47, PM = 3.03, 95% CrI = 0.24 to 0.76, P_MCMC_ = 0.025, Supplementary Fig 2). Bayesian pairwise contrasts further indicated that females had lower hearing thresholds than males at 1 kHz (P_MCMC_ = 0.025), 2 kHz (P_MCMC_ = 0.0006), 4 kHz (P_MCMC_ = 0.002), 8 kHz (P_MCMC_ = 0.021), and16 kHz (P_MCMC_ < 0.001). However, males and females exhibited no significant difference in hearing thresholds at 24 kHz (P_MCMC_ = 0.072), 32 kHz (P_MCMC_ = 0.528), 46 kHz (P_MCMC_ = 0.144), and 64 kHz (P_MCMC_ = 0.248).

When examining sex differences within social categories (Fig 2C), PGLMM analyses revealed no significant differences in auditory evoked thresholds between solitary males and solitary females across all tested frequencies (PGLMM: H^2^ = 0.16, PM = 2.57, 95% CrI = -0.79 to 6.37, P_MCMC_ = 0.152, Fig 2C). Similarly, no significant differences in auditory thresholds were detected between social males and social females across tested frequencies (PGLMM: H^2^ = 0.16, PM = 3.24, 95% CrI = -0.97 to 7.14, P_MCMC_ = 0.116, Fig 2C). However, when sex specific variation was examined between social categories, PGLMM analyses revealed that solitary males exhibited higher auditory evoked thresholds than social males (PGLMM: H^2^ = 0.16, PM = 11.51, 95% CrI = 6.08 to 17.07, P_MCMC_ = 0.002) and social females when considering all tested frequencies (PGLMM: H^2^ = 0.16, PM = 14.75, 95% CrI = 9.12 to 19.87, P_MCMC_ = 0.002). Bayesian pairwise contrasts further indicated that solitary males consistently exhibited higher auditory thresholds than social females across all tested frequencies (All P_MCMC_ < 0.05, Supplementary file 1). Solitary males also displayed higher auditory thresholds than social males across most tested frequencies (1-24 kHz, All P_MCMC_ < 0.05), except at 32, 46, and 64 kHz, where both groups exhibited no difference in auditory thresholds (Supplementary file 1). Similarly, solitary females generally exhibited higher auditory thresholds than both social males (PGLMM: H^2^ = 0.16, PM = 8.94, 95% CrI = 4.05 to 13.9, P_MCMC_ = 0.004) and social females (PGLMM: H^2^ = 0.16, PM = 12.18, 95% CrI = 6.72 to 17.27, P_MCMC_ = 0.002). Pairwise analyses further revealed that solitary females exhibited higher auditory thresholds than social males at 1kHz (P_MCMC_ =0.004), 2 kHz (P_MCMC_ = 0.004), 4 kHz (P_MCMC_ = 0.004), and 8 kHz (P_MCMC_ = 0.002), but not at the remaining tested frequencies (16-46 kHz, P_MCMC_ > 0.05). Solitary females also displayed higher auditory thresholds than social females across all tested frequencies (P_MCMC_ < 0.05), except at 64 kHz (P_MCMC_ = 0.072).

### ABR Amplitude Ratios

Amplitude ratios were calculated as the ratio of ABR peak wave IV amplitude to peak wave I amplitude for the left and right pinnae at each intensity (50 – 90 dB SPL). This metric reflects the strength of neural activity relative to earlier response components and provides a normalized measure (which accounts for species variation in body size) of auditory signal transmission through the ascending auditory system. In both sexes, amplitude ratios increased monotonically with increasing intensity (50 – 90 dB SPL). When analyzing the data for sex differences across all species, independent of sociality, PGLMM analyses revealed significantly higher amplitude ratios in males than females across tested intensities (PGLMM: H^2^ = 0.03, PM = 0.25, 95% CrI = 0.17 to 0.33, P_MCMC_ < 0.001, Supplementary Fig 3A). Bayesian pairwise contrasts further indicated that males had higher amplitude ratios than females at 50 dB SPL (P_MCMC_ < 0.001) and 60 dB SPL (P_MCMC_ < 0.001), whereas no sex differences were detected at higher intensities (70 dB SPL: P_MCMC_ = 0.944; 80 dB SPL: P_MCMC_ = 0.736; 90 dB SPL: P_MCMC_ = 0.088).

When social lifestyle strategies were considered to assess sex differences within groups, PGLMM analyses revealed significant differences in amplitude ratios between solitary males and solitary females (PGLMM: H^2^ = 0.04, PM = 0.18, 95% CrI = 0.05 to 0.29, P_MCMC_ = 0.008, Fig 3A). Bayesian pairwise contrasts indicated that solitary males had higher amplitude ratios than solitary females at 50 dB SPL (P_MCMC_ = 0.008), whereas no differences were detected at 60 dB SPL (P_MCMC_ = 0.260), 70 dB SPL (P_MCMC_ = 0.310), 80 dB SPL (P_MCMC_ = 0.350), or 90 dB SPL (P_MCMC_ = 0.050). Similarly, significant sex differences were also detected between social males and social females, with social male rodents exhibiting higher amplitude ratios overall (PGLMM: H^2^ = 0.04, PM = 0.32, 95% CrI = 0.21 to 0.42, P_MCMC_ < 0.001). Bayesian pairwise contrasts further showed that social males had higher amplitude ratios than social females at 50 dB SPL (P_MCMC_ <0.001) and 60 dB SPL (P_MCMC_ <0.001), but not at higher intensities (70-90 dB SPL, all P_MCMC_ > 0.05, Supplementary file 2). When testing for sex differences between social categories, PGLMM further revealed significant differences in amplitude ratios between solitary males and social females (PGLMM: H^2^ = 0.04, PM = 0.2, 95% CrI = 0.07 to 0.32, P_MCMC_ = 0.002). Solitary males exhibited higher amplitude ratios than social females at 50 dB SPL (P_MCMC_ = 0.002), and 60 dB SPL (P_MCMC_ = 0.040), but not at 70 dB SPL (P_MCMC_ = 0.914), 80 dB SPL (P_MCMC_ = 0.450), or 90 dB SPL (P_MCMC_ = 0.090). PGLMM analyses further revealed significant sex differences in amplitude ratios between solitary females and social males (PGLMM: H^2^ = 0.04, PM = 0.02, 95% CrI = 0.07 to 0.32, P_MCMC_ = 0.002, Fig 3A). Overall, solitary females exhibited lower amplitude ratios than social males at 50 dB SPL (P_MCMC_ = 0.002) and 60 dB SPL (P_MCMC_ = 0.040), whereas no differences were detected at 70 dB SPL (P_MCMC_ = 0.914), 80 dB SPL (P_MCMC_ = 0.450), or 90 dB SPL (P_MCMC_ = 0.090). In contrast, no significant differences in amplitude ratios were detected between solitary males and social males across tested intensities (PGLMM: H^2^ = 0.04, PM = -0.12, 95% CrI = -0.07 to 0.32, P_MCMC_ = 0.074). Likewise, no significant differences in amplitude ratios were detected between solitary females and social females across tested intensities (PGLMM: H^2^ = 0.04, PM = 0.02, 95% CrI = -0.09 to 0.14, P_MCMC_ = 0.734).

**Fig 3A:**
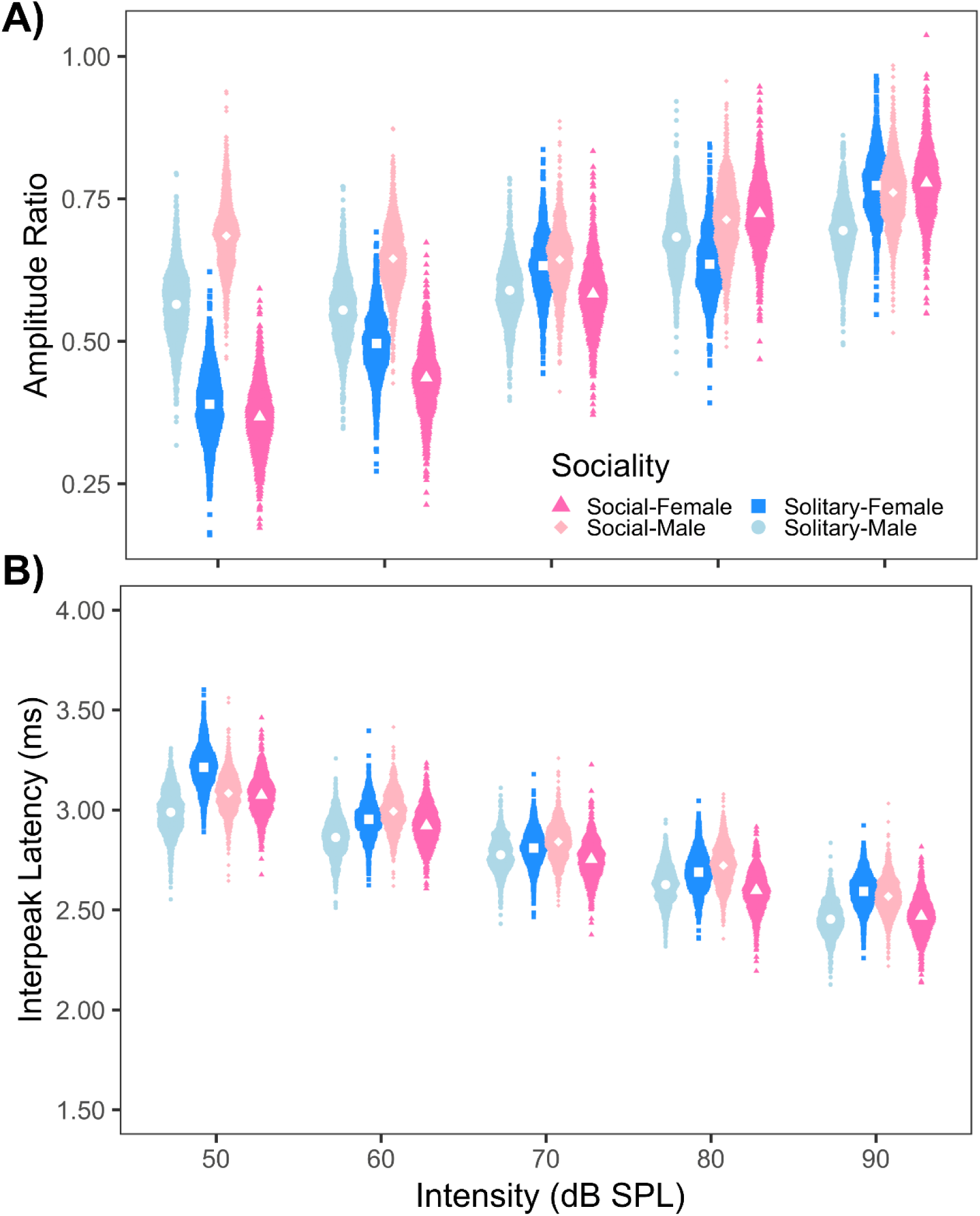
Average amplitude ratios between the sexes across social groups. **B** shows Average interpeak latencies across intensities [(n = 180, 20 individuals per species (10 males and 10 females)]. Significant sex differences were detected for amplitude ratios, but not for interpeak latencies across social categories.

### ABR Interpeak Latencies

Interpeak latencies were calculated as the difference between the peak latencies of wave I and IV at each intensity for both left and right monoaural responses. This metric served as an index of neural conduction time in the ascending auditory system. As expected, interpeak latencies decreased as intensity increased in both sexes. The results of the PGLMM model revealed no overall sex differences in wave I-IV interpeak latencies across species when all species were analyzed independent of sociality (PGLMM: H^2^ = 0.03, PM = -0.09, 95% CrI = - 0.23 to 0.05, P_MCMC_ = 0.210, Supplementary Fig 3B).

Similarly, comparisons within and between social categories revealed no significant sex differences in interpeak latencies. Specifically, no significant differences were detected between solitary males and solitary females (PGLMM: H^2^ = 0.03, PM = -0.23, 95% CrI = - 0.46 to 0.01, P_MCMC_ = 0.054, Fig 3B), or between social males and social females (PGLMM: H^2^ = 0.02, PM = 0.01, 95% CrI = - 0.18 to 0.21, P_MCMC_ = 0.920, Fig 3B). Likewise, no significant differences were detected among any cross-category comparisons, including solitary males and social males (PGLMM: H^2^ = 0.03, PM = -0.1, 95% CrI = - 0.33 to 0.13, P_MCMC_ = 0.462), solitary males and social females (PGLMM: H^2^ = 0.03, PM = -0.09, 95% CrI = - 0.3 to 0.14, P_MCMC_ = 0.468), solitary females and social males (PGLMM: H^2^ = 0.03, PM = 0.13, 95% CrI = - 0.09 to 0.35, P_MCMC_ = 0.242), or solitary females and social females (PGLMM: H^2^ = 0.03, PM = 0.14, 95% CrI = - 0.08 to 0.36, P_MCMC_ = 0.200, Supplementary file 3). Because there were no sex-differences in interpeak latencies across any social categories, Bayesian pairwise contrast tests were not performed.

### BIC Relative Amplitudes

BIC ABR peak amplitudes and latencies were extracted at each ITD to assess for sex differences in binaural hearing sensitivity across tested ITDs. When data were pooled across species independent of sociality, PGLMM analyses revealed significant sex differences in BIC relative amplitudes across ITDs (PGLMM: H^2^ = 0.11, PM = -6.91, 95% CrI = -11.95 to -1.89, P_MCMC_ = 0.004, Supplementary Fig 4A). Bayesian pairwise contrasts further indicated that females exhibited higher relative amplitude than males at 0.5 ms ITD (P_MCMC_ = 0.004). In contrast, no differences were detected between the sexes at 1.0 ms (P_MCMC_ = 0.708), 1.5 ms (P_MCMC_ = 0.261), and 2.0 ms ITDs (P_MCMC_ = 0.260).

When testing for sex differences within social categories, phylogenetic comparative analyses further revealed significant differences in BIC relative amplitudes between solitary males and solitary females (PGLMM: H^2^ = 0.11, PM = - 8.27, 95% CrI = - 15.72 to – 1.16, P_MCMC_ = 0.018, Fig 4A). Bayesian pairwise contrasts indicated that solitary females exhibited higher BIC relative amplitudes than solitary males at 0.5 ms ITDs (P_MCMC_ = 0.018), but not at 1.0 ms (P_MCMC_ = 0.364), 1.5 ms (P_MCMC_ = 0.940) or 2.0 ms ITDs (P_MCMC_ = 0.852). In contrast, no significant sex differences in BIC relative amplitudes were detected between social males and social females across tested ITDs (PGLMM: H^2^ = 0.11, PM = - 5.72, 95% CrI = - 12.86 to 1.4, P_MCMC_ = 0.122, Supplementary file 4). When sex differences were assessed between social categories, PGLMM revealed significant differences in BIC relative amplitudes between solitary males and social females (PGLMM: H^2^ = 0.11, PM = - 12.8, 95% CrI = - 21.18 to – 3.62, P_MCMC_ = 0.004). Social females exhibited higher BIC relative amplitudes than solitary males at 0.5 ms ITDs (P_MCMC_ = 0.004), whereas no differences were detected at 1.0 ms (P_MCMC_ = 0.150), 1.5 ms (P_MCMC_ = 0.182) or 2.0 ms ITDs (P_MCMC_ = 0.466). No significant differences in BIC relative amplitudes were detected between solitary males and social males across tested ITDs (PGLMM: H^2^ = 0.11, PM = - 7.08, 95% CrI = - 15.82 to – 2.02, P_MCMC_ = 0.116). Likewise, no overall differences in BIC relative amplitudes were detected between solitary females and social males (PGLMM: H^2^ = 0.11, PM = 1.19, 95% CrI = - 7.35 to – 10.68, P_MCMC_ = 0.812) or between solitary females and social females across tested ITDs (PGLMM: H^2^ = 0.11, PM = - 4.53, 95% CrI = - 12.77 to 4.48, P_MCMC_ = 0.308).

**Fig 4A:**
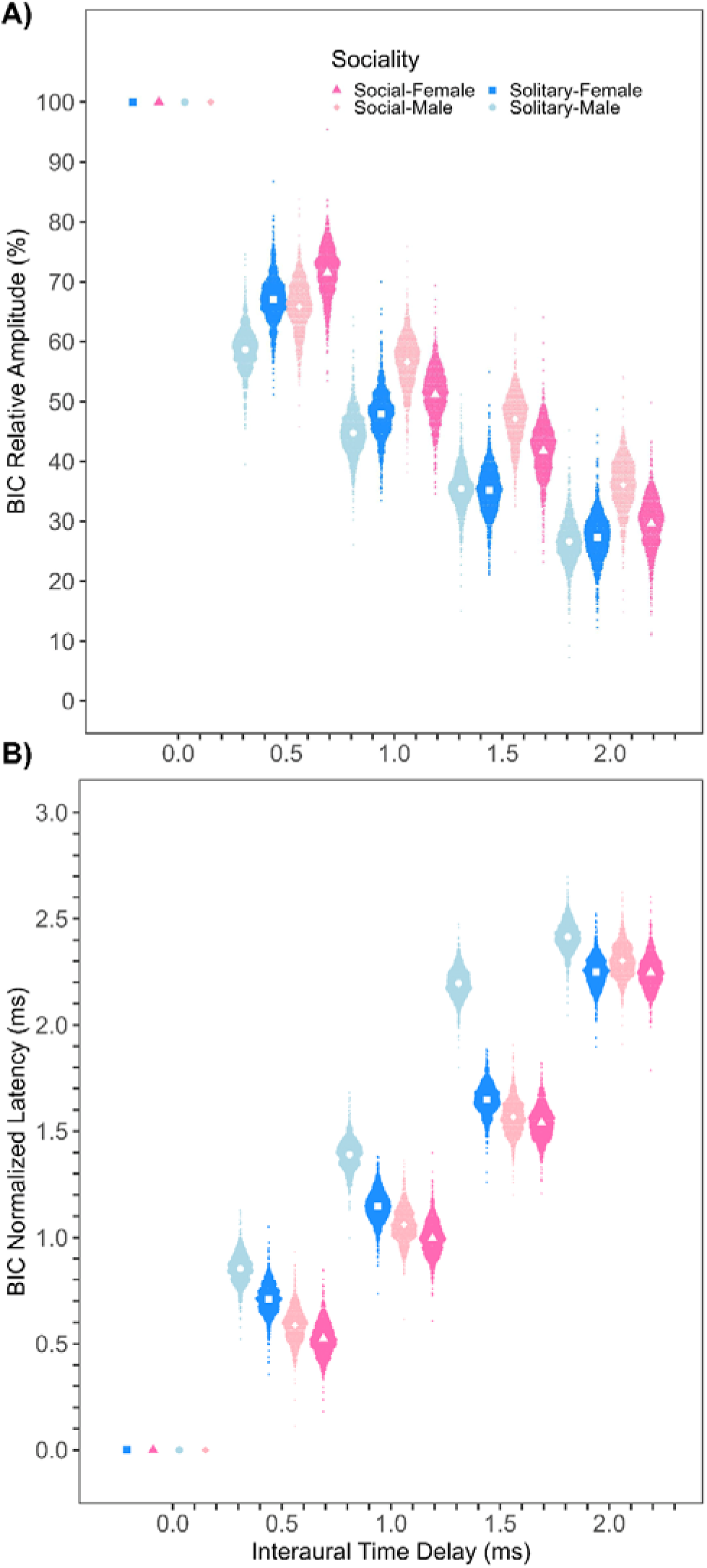
Average BIC relative amplitude between the sexes across social categories. **B** shows Average BIC normalized latencies across ITDs [(n = 180, 20 individuals per species (10 males and 10 females)]. Significant sex differences were detected in both BIC relative amplitude and BIC normalized latencies across social categories.

### BIC Normalized Latencies

When all species were analyzed independent of sociality to test for sex differences, phylogenetic comparative analyses revealed substantial differences in BIC normalized latencies between female and male rodents across tested ITDs (PGLMM: H^2^ = 0.12, PM = 0.10, 95% CrI = 0.006 to 0.21, P_MCMC_ = 0.040, Supplementary Fig 4B). Pairwise contrasts indicated that females exhibited faster BIC normalized latencies than males at 0.5 ms (P_MCMC_ = 0.040), 1.0 ms (P_MCMC_ = 0.004), and 1.5 ms ITDs (P_MCMC_ < 0.001), but not at 2.0 ms ITDs (P_MCMC_ = 0.064).

When analyses were conducted for sex differences within social categories, significant differences in BIC normalized latencies were detected between solitary males and solitary females (PGLMM: H^2^ = 0.03, PM = 0.15, 95% CrI = 0.0 to 0.3, P_MCMC_ = 0.044, Fig 4B). Bayesian pairwise contrasts further indicated that solitary females exhibited faster BIC normalized latencies than solitary males at 0.5 ms (P_MCMC_ = 0.044), 1.0 ms (P_MCMC_ < 0.001), 1.5 ms (P_MCMC_ < 0.001), and 2.0 ms ITDs (P_MCMC_ = 0.020). However, no significant differences were detected in BIC normalized latencies between social males and social females (PGLMM: H^2^ = 0.03, PM = 0.06, 95% CrI = -0.09 to 0.2, P_MCMC_ = 0.426, Supplementary file 5). For analyses of sex differences between social categories, PGLMM revealed significant sex differences in BIC normalized latencies between solitary males and social males (PGLMM: H^2^ = 0.03, PM = 0.27, 95% CrI = 0.1 to 0.43, P_MCMC_ = 0.002), and between solitary males and social females (PGLMM: H^2^ = 0.03, PM = 0.33, 95% CrI = 0.17 to 0.49, P_MCMC_ = 0.002). Social males exhibited faster BIC normalized latencies that solitary males at 0.5 ms (P_MCMC_ = 0.002), 1.0 ms (P_MCMC_ = 0.002), 1.5 ms ITDs (P_MCMC_ < 0.001), but not at 2.0 ms ITDs (P_MCMC_ = 0.180). Social females also exhibited faster BIC normalized latencies than solitary males at 0.5 ms (P_MCMC_ = 0.002), 1.0 ms (P_MCMC_ < 0.001), 1.5 ms (P_MCMC_ < 0.001), and 2.0 ms (P_MCMC_ = 0.046). In addition, PGLMM analyses identified significant differences in BIC normalized latencies between solitary females and social females (PGLMM: H^2^ = 0.03, PM = 0.18, 95% CrI = 0.01 to 0.35, P_MCMC_ = 0.040). Social females exhibited faster BIC normalized latencies than solitary females at 0.5 ms ITDs (P_MCMC_ = 0.040), but not at 1.0 ms (P_MCMC_ = 0.090), 1.5 ms (P_MCMC_ = 0.226), and 2.0 ms ITDs (P_MCMC_ = 0.986). No significant sex differences in BIC normalized latencies were detected between solitary females and social males (PGLMM: H^2^ = 0.03, PM = 0.12, 95% CrI = -0.05 to 0.3, P_MCMC_ = 0.162).

## DISCUSSION

The central goal of this study was to examine whether sociality drives sex differences in auditory processing across wild-caught rodents. To address this question, we assessed sex-specific variations in frequency and click-evoked thresholds as measures of hearing sensitivity, as well as monaural and binaural ABR amplitudes and latencies, which reflect peripheral and central neural activity and binaural auditory sensitivity within the ascending auditory pathway. Across social groups, females consistently exhibited enhanced auditory sensitivity relative to males, including lower click and frequency evoked thresholds, higher BIC relative amplitude, and faster BIC normalized latency, patterns indicative of increased neural sensitivity. In combination with low phylogenetic signals detected across auditory traits, these findings suggest that enhanced auditory sensitivity in females may represent a broadly conserved characteristic of rodent auditory processing rather than a pattern strongly constrained by phylogenetic relatedness. Collectively, these findings demonstrated that sex is an important source of variation in rodent auditory function and may interact with ecological, behavioral, and social selective pressures to shape auditory system organization of the studied rodents included in this investigation.

Our phylogenetic comparative analyses revealed that both sexes exhibited the greatest hearing sensitivity between 8 and 24 kHz, with females consistently showing lower auditory thresholds than males at 1, 2, 4, 8, and 16 kHz when data were pooled across species, independent of sociality. Similarly, substantial sex differences in click thresholds were detected across social categories, except for solitary species, where males and females showed comparable click thresholds. Importantly, sex-specific differences in hearing sensitivity at specific frequencies were more pronounced in social than solitary rodents, indicating that sociality may amplify auditory divergence between the sexes across rodent species reported here. Across social groups, social rodents exhibited auditory thresholds approximately 10-15 dB SPL lower than solitary rodents. For example, auditory thresholds at 8 and 16 kHz were about 10 dB SPL higher in solitary males than in social males; however, the disparity increased to approximately 15 dB SPL when comparing social females with solitary males. These findings suggest that sociality is an important driver of auditory trait divergence and may shape the evolution of sex-specific hearing sensitivity in the nine rodent species measured in the current study.

In addition, the enhanced hearing sensitivity observed in females is consistent with previous studies in laboratory rodents and humans, where females often exhibit lower auditory thresholds than males^25,28, 58–61^. Such sex differences in auditory function have been attributed to a combination of factors, including cochlear mechanics, neural synchrony, and hormonal modulation of auditory pathways^24,27,79–82^. However, much of the existing literature has focused mainly on domesticated laboratory rodents raised under controlled acoustic and social conditions, raising questions regarding the extent to which these findings generalize to natural populations. By demonstrating similar patterns across wild-caught rodents spanning diverse social systems and evolutionary histories, our findings extend previous work and suggest that the enhanced female sensitivity to quieter sounds represents a robust biological phenomenon rather than an artifact of laboratory settings. Moreover, the persistence and amplification of these sex differences among social groups, despite substantial interspecific variation and low phylogenetic signals, indicate that sex-related auditory advantages may arise through conserved physiological mechanisms or repeated convergent evolution in response to shared ecological, communication or parental care demands among the species examined here. Collectively, these findings support our first hypothesis that social species would exhibit lower frequency response thresholds within rodents’ peak hearing range, as well as exhibit lower click thresholds, relative to solitary species.

Rodents rely extensively on ultrasonic vocalizations (USVs) to mediate social interactions, and previous studies have documented pronounced sex differences in the structure and context of USVs across species with varying social structures^83,84^. In social rodents, females generally produce fewer USVs and emit calls at slightly higher frequencies than males^85–88^, whereas solitary species, often exhibit limited or no sex differences in call duration, frequency, or call rate^89^. Additionally, rodents are known to produce sonic and ultrasonic calls ranging from 1 to 110 kHz, with peak energy most commonly concentrated between 10 and 35 kHz^25,86,90^. However, these frequency ranges are based on studies of select rodent taxa, and species-specific vocalization data are not available for all species examined here. While all species demonstrated auditory sensitivity across the frequency spectrum of rodent vocalizations, both sexes exhibited peak hearing sensitivity between 8 and 24 kHz, closely aligning with the dominant spectral components of rodents’ typical calls, consistent with predictions of the matched filter hypothesis^91^. Particularly, the larger sex differences in hearing sensitivity observed in social species suggest that social organization may shape auditory turning in ways that enhance the detection of behaviorally relevant acoustic cues and offspring-related vocal signals. In addition, lower frequency sounds propagate more effectively over long distances, and social species such as *Mus musculus*, *Onychomys leucogaster* and *Peromyscus californicus* have been reported to produce relatively low-frequency calls when separated from conspecifics^85,92^. Enhanced low-frequency hearing sensitivity in females may therefore improve long-distance communication and social cohesion under natural environmental conditions. More broadly, these findings support the idea that sociality contributes to the evolution of sexual dimorphism in auditory traits across rodents by shaping the selective pressure acting on acoustic communication systems. Future comparative studies across tetrapods using ecologically relevant acoustic stimuli and naturalistic social contexts will be important for clarifying the functional significance of enhanced low-frequency sound detection in females across different social structures and environmental conditions.

Amplitude ratios increased with stimulus intensity in both sexes across sociality categories, with stronger neural gain within the ascending auditory pathway at higher sound levels, consistent with previous studies^24,60,67^. However, sex differences in amplitude ratios varied with sociality, suggesting that social organization contributes to sexually dimorphic auditory processing in species reported here. When data were analyzed independent of sociality, males exhibited higher amplitude ratios than females at lower intensities (50-60 dB SPL), but no differences were detected at higher intensities (70-90 dB SPL). When analyzing across social groups, these differences were also most pronounced in solitary species, where solitary males showed higher amplitude ratios than solitary and social females. In addition, no overall differences were detected between solitary and social females or between solitary and social males. Higher amplitude ratios in solitary males may therefore reflect stronger early auditory nerve activity or increased peripheral neural gain, consistent with previous reports^60,67^. Accordingly, the larger sex differences observed among social systems further suggest that social communication may shape auditory processing differently in male and female rodents. In socially complex environments, males may benefit from enhanced encoding of low-intensity acoustic cues associated with territorial defense or mate competition, while females may exhibit adaptations favoring increased auditory sensitivity such as lower auditory thresholds. Together, these findings support the hypothesis that sociality drives the evolution of sexual dimorphic auditory traits in rodents by amplifying males’ auditory neural gains over females.

Consistent with previous studies^24,60,67^, interpeak latencies decreased with increasing stimulus intensity in both sexes. In contrast to amplitude ratios, no overall significant sex differences in interpeak latencies were detected either across all species or between sociality categories. These findings indicated that, although sociality shapes sexual dimorphic variation in auditory response amplitude, it has no effect on neural conduction timing within the ascending auditory system of these studied species. The lack of sociality-related differences in interpeak latencies suggests that neural conduction timing may be more evolutionary conserved than amplitude-based measures of auditory processing. Therefore, different components of the auditory system may evolve under distinct selective pressures, with amplitude-related traits appearing more responsive to solitary selective forces than temporal conduction properties. Collectively, these results demonstrate that sociality contributes to the evolution of sexual dimorphism in rodent auditory processing, particularly through the effects on neural response amplitude rather than conduction timing.

Our findings showed that BIC relative amplitude decreased with increasing ITD, a pattern consistent with previous comparative studies of mammalian binaural auditory processing^24,29,60,63,68^. Across all species, females exhibited significantly greater BIC relative amplitudes than males, particularly at 0.5 ms ITDs, suggesting enhanced neural encoding of binaural stimuli at short interaural delays. When sociality was incorporated into the analyses, similar patterns were observed among most social systems. Particularly, social females exhibited significantly greater BIC relative amplitude than solitary males. The elevated BIC relative amplitude sensitivity observed in social females relative to solitary males, may reflect enhanced neural encoding of interaural cues important for sound localization and communication. In natural grassland habitats, high frequency vocal signals attenuate more rapidly than low-frequency signals, which can limit the distance over which high frequency sounds are detected. As all species were captured in grassland environments, enhanced BIC amplitudes in females of social rodents may therefore facilitate the detection and localization of socially relevant vocalizations under acoustically challenging conditions. Because differences in BIC amplitudes were largest between solitary males and social females, but largely absent between the sexes in social species, our results suggest that sociality shapes the magnitude and direction of sex differences in binaural processing rather than sex alone independently determining auditory performance. Our findings further suggest that sex differences in binaural processing may be functionally tuned to ecological and social demands, particularly in species where precise sound localization confers fitness advantages.

In accordance with previous studies^29,60,63,68^, BIC normalized latency decreased with increasing ITD, and females generally exhibited faster BIC normalized latencies than males. Across all species, females showed significantly faster BIC normalized latencies than males at 0.5 ms, 1.0 ms and 1.5 ms ITD, indicating relatively faster central processing of binaural information within auditory pathways involved in sound localization^63,68^. Faster neural timing may allow females to decode spatial information from vocal signals more efficiently, thereby improving communication, coordination of social interactions, and rapid responses to environmental acoustic cues. Importantly, the inclusion of sociality revealed that sex differences in binaural temporal processing were shaped by social lifestyle strategy. Solitary males consistently exhibited slower normalized latencies than social males, and social females at most ITDs, while differences among the other groups were limited or absent. For instance, social females exhibited faster latencies than solitary males across all tested ITDs, suggesting that social living may further enhance rapid binaural processing in females. In contrast, no significant differences were detected between solitary females and social males or between social males and social females, indicating convergence in binaural timing among socially integrated groups. Together, these patterns suggest that sociality is a major driver of sex differences in auditory processing in rodents. Rather than reflecting fixed sex-based auditory specialization, binaural temporal processing appears to vary according to the communication and ecological demands associated with different social systems. Solitary males may experience reduced selective pressure for rapid binaural processing compared with females and socially living individuals, whereas females and social species may benefit from enhanced temporal precision for detecting and localizing socially relevant vocalization. The smaller sex differences observed within social groups further support the idea that frequent social interactions and shared acoustic environment may promote convergence in auditory processing between males and females.

Although ABR measures do not directly assess auditory perception or behavior, they are widely used as physiological markers of auditory sensitivity and brainstem temporal processing across taxa. Accordingly, variation in ABR thresholds and binaural auditory responses observed between the sexes and across social groups may reflect differences in the early neural encoding of auditory information. Comparative studies across taxa further suggest that auditory sensitivity can evolve alongside social complexity, potentially because individuals in larger or more socially interactive groups must detect and discriminate a greater number of calls and socially relevant vocal signals across heterogeneous environments^10^. For instance, research in primates has demonstrated that species forming larger social groups tend to exhibit enhanced hearing sensitivity, consistent with a social drive hypothesis in which increasing sociality selects for improved auditory detection and processing of communication signals^10^. Similarly, studies in zebra finches have reported correlations between measures of vocal responsiveness and neural responses properties during the perception of familiar calls. Individuals exhibited higher vocal responses rates, faster response latencies, stronger and more prolonged neural activity in auditory-responsive forebrain regions^93^. These results suggest a potential association between the neural encoding of socially relevant vocal signals and behavioral responsiveness, providing further support for the hypothesis that auditory processing and social communication may co-evolve. Nevertheless, similar to previous comparative work, the present findings do not establish a causal link between auditory physiology and behavioral performance. Because auditory perception and communication behavior were not directly measured, these interpretations should be regarded as hypotheses regarding potential functional implications rather than demonstrated behavioral outcomes. Future studies incorporating neurophysiological measurements with behavioral essays of sound detection, localization, or acoustic communication will be necessary to determine whether auditory differences observed here correspond to ecologically meaningful variation in sensory performance between the sexes among rodents with different social lifestyle strategies.

While our study shows sex-differences in auditory brainstem responses among social groups, direct comparisons with previously published literature can be challenging. One limitation of the current work is the use of anesthesia during ABR recordings. Previous research has shown that ABR thresholds measured under anesthesia are typically 10-20 dB higher than those recorded using behavioral methods^94,95^. Therefore, the auditory thresholds reported here may overestimate true behavioral sensitivity. Future studies incorporating behavioral assessment will be essential to directly confirm the results reported here. In addition, all individuals tested in this study were wild-caught, and precise age and reproductive status, particularly of females were unknown. These factors are known to influence auditory processing and could contribute to interindividual variability^24,79–82^. Further work that controls for age, hormonal status, and reproductive condition will be necessary for refining interpretations of sex-specific auditory differences. Finally, auditory thresholds in this study were determined through visual inspection, which may introduce subjectivity^96^. However, prior studies have demonstrated minimal differences between observer-based methods and automated algorithms for threshold detection^24,29,60,67,97^. Nonetheless, further validation of our observer method with more quantitative algorithms would be essential to confirm threshold values reported here, although our thresholds are in well accordance with previously published work, giving us confidence in the findings reported in this present investigation.

## CONCLUSION

Our results reveal consistent sex-specific differences in auditory processing across wild-caught rodent species representing distinct social lifestyle organization. We showed that both sexes exhibited click and frequency-specific variation in auditory sensitivity, with peak hearing sensitivity occurring between 8 and 24 kHz. Males generally exhibited higher ABR amplitude ratios than females, whereas interpeak latencies were largely conserved between the sexes across social groups. In addition, we showed that females generally exhibited higher BIC relative amplitude and faster BIC normalized latency than males at most tested ITDs. However, these sex-differences were influenced by sociality. We demonstrated that auditory processing in the studied rodents reported here reflects an interaction between sex and sociality, likely driven by differences in communication demands and ecological pressures associated with distinct social systems. Future behavioral, neurophysiological, and comparative studies across a broader range of vertebrates will be essential to determine whether these patterns reflect evolutionarily conserved auditory strategies among tetrapods and to clarify how social behavior influences the evolution of sensory systems.

## ACKNOWLEGEMENTS

We thank the personnels of the Packsaddle, James Collin, and Sandy Sanders wildlife management areas, Sheena Parsons, Benny Farrar, Marie Stone, and Marcus Thibodeau for housing and permission to collect rodents in our sampling locations. We also thank all undergraduates on team wild rodent of the McCullagh lab for helping in trapping to complete this manuscript. We also thank Dr. Tim Lei and Dr. Ben-Zheng Li for developing the custom ABR software and analysis program. This work was partially supported by the Payne County Audubon Society through the Helen Miller research grant, which helped in covering fieldwork costs.

## SUPPLEMENTARY MATERIALS AND DATA ACCESSIBIBILITY

Supplementary materials and all raw data required to reproduce the figures and analyses of this current investigation is available online at: ADD link here please

## AUTHORS’ CONTRIBUTIONS

L.J: Conceptualization, data analysis, investigation, methodology, validation, visualization, writing-original draft

D.M.J: Conceptualization, data analysis, investigation, validation, methodology, visualization

N.C.R: Conceptualization, investigation, validation, methodology, visualization

F.A.M: Conceptualization, validation, data analysis, visualization

E.A.M: Conceptualization, data analysis, investigation, methodology, validation, visualization

## CONFLICT OF INTEREST DECLARATION

We declare we have no competing interests.

## FUNDING

No sources of fundings were associated with this work.

## Notes

### Competing Interest Statement

The authors have declared no competing interest.

