## supplemental figures and tables for "Sociality Drives Sex Differences in Auditory Brainstem Processing Across Rodent Species"

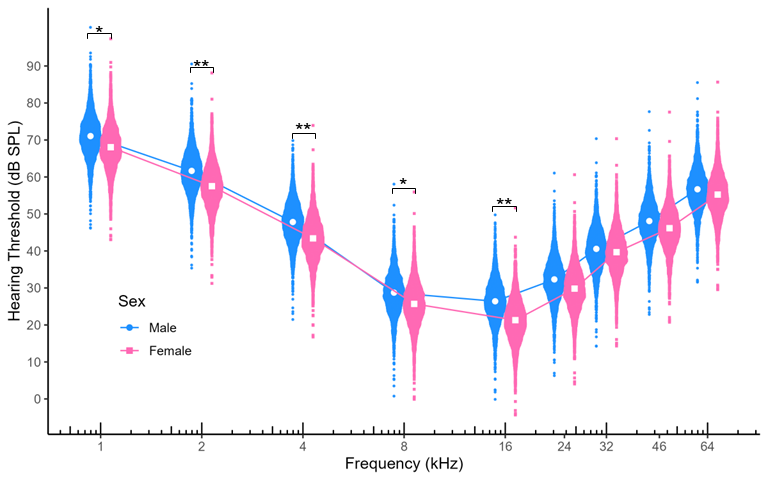

**Fig S1:** Mean auditory brainstem response thresholds between male and females wild-caught rodents across tested frequencies (n = 180, 20 individuals per species (10 males and 10 females). Colors represent sexes: blue = male, and pink = females. Significant differences were detected in frequency evoked response thresholds between sexes. Asterisks represent Bayesian statistical differences between sexes with p < 0.05 *, p ≤ 0.01 **.

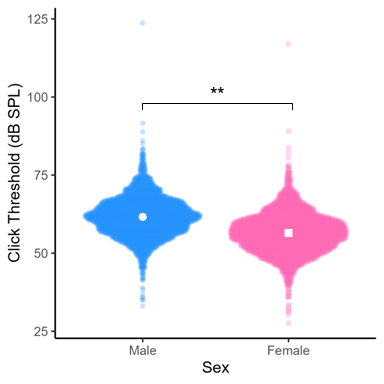

**Fig S2:** Mean click thresholds between male and females wild-caught rodents across tested frequencies (n = 180, 20 individuals per species (10 males and 10 females). Colors represent sexes: blue = male, and pink = females. Significant differences were detected in click thresholds between sexes. Asterisks represent Bayesian statistical differences with p < 0.01 **.

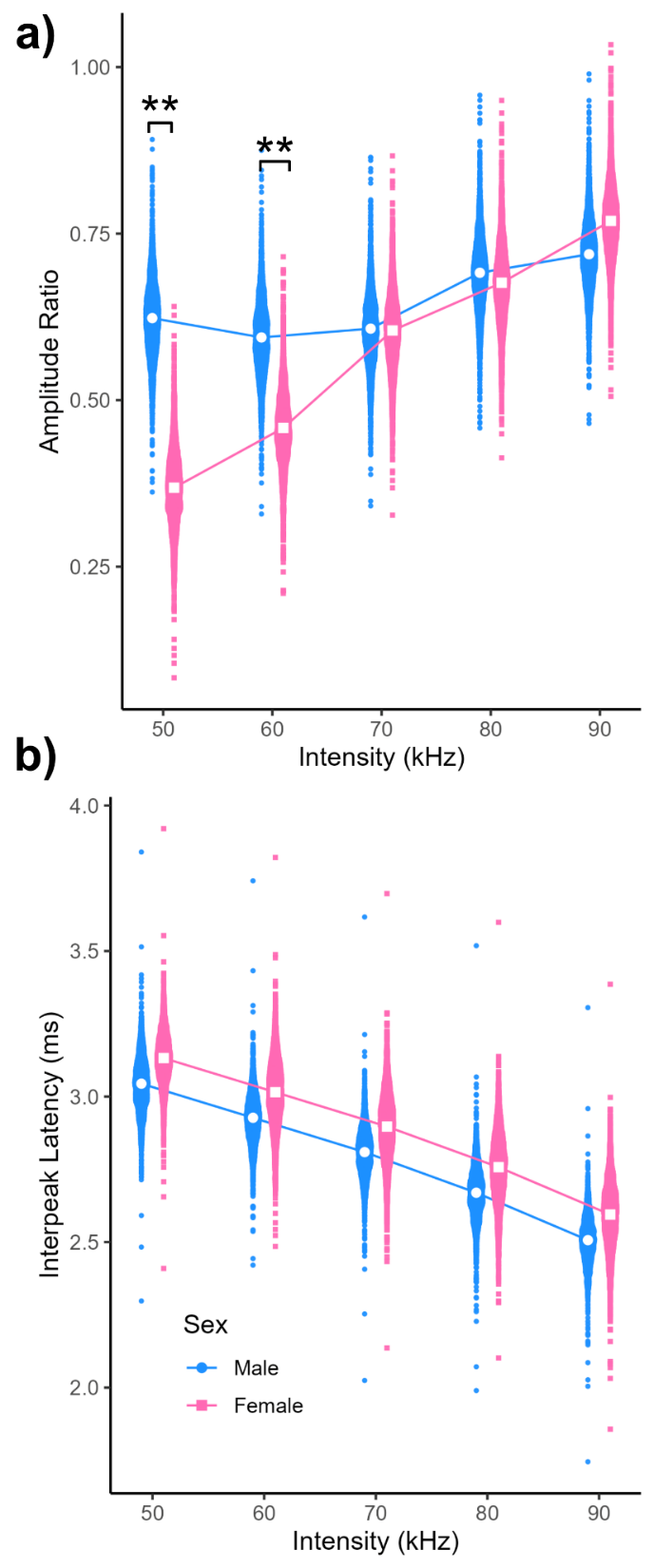

**Fig S3:** **a**: Average amplitude ratio of ABR wave I and IV between the sexes (n = 180, 20 individuals per species (10 males and 10 females). **b** represent average interpeak latency of ABR wave I and IV between the sexes across intensities (n = 180, 20 individuals per species (10 males and 10 females). Colors represent sexes: blue = male, pink = female. Significant differences were detected in amplitude ratio and interpeak latency between the sexes. Asterisks represent Bayesian statistical differences between the sexes with p < 0.01 **.

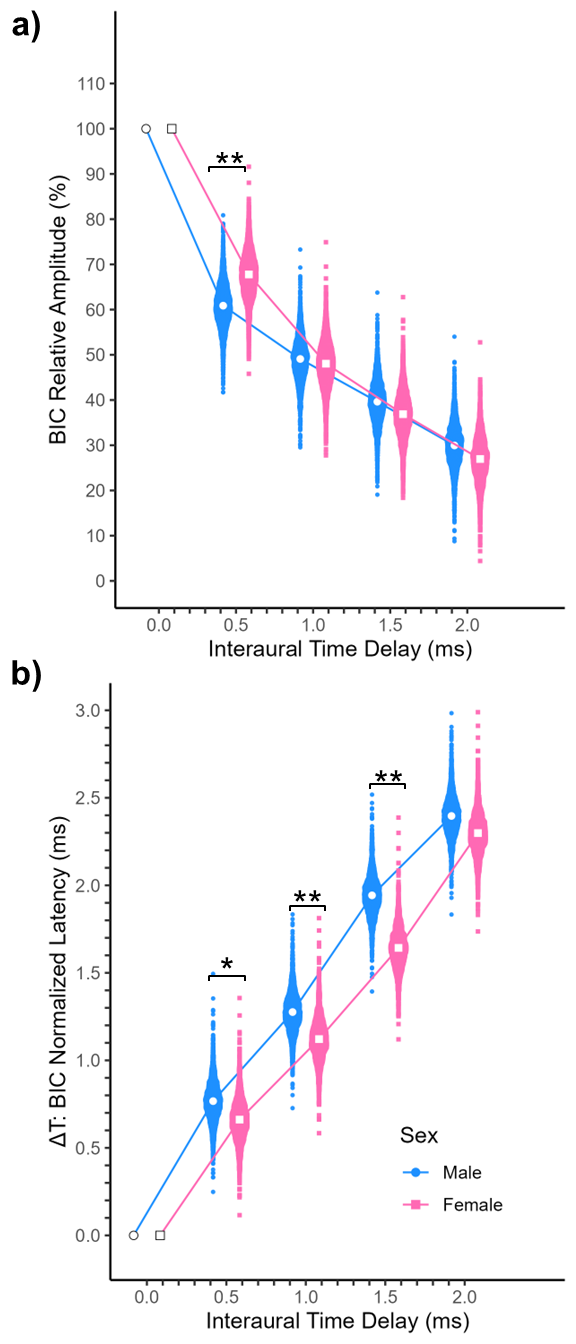

**Fig S4:** **a)** Average BIC relative amplitude between the sexes across ITD (n = 180, 20 individuals per species (10 males and 10 females). **b** represent average normalized latency between the sexes across ITD (n = 180, 20 individuals per species (10 males and 10 females). Colors represent sexes: blue = male, pink = female. Significant differences were detected in relative amplitude and normalized latency between the sexes. Asterisks represent Bayesian statistical differences between the sexes with p < 0.05 *, p ≤ 0.01 **.

**Table S1:** Deviance Information Criterion (DIC) of all models

| **Frequency Evoked Threshold** | | | |
| --- | --- | --- | --- |
| **Models** | **Deviance Information Criterion (DIC)** | **Best Model** | |
| **Null:** dB ~ Frequency | 11349.30 |  | |
| **Additive:** dB ~ Sociality + Sex + Frequency | 11347.31 |  | |
| **Full:** dB ~ Sociality * Sex * Frequency | 11284.54 |  | |
| **Full + BM:** dB ~ Sociality * Sex * Frequency + BM | 11285.63 |  | |
| **Click Threshold** | | | |
| **Models** | **Deviance Information Criterion (DIC)** | | **Best Model** |
| **Null:** dB ~ 1  **Additive: dB ~** Sociality + Sex | 1098.33  1035.11 | | |
| **Full: dB** ~ Sociality * Sex | 1025.67 | |  |
| **Full + BM:** dB ~ Sociality * Sex + BM | 11285.69 | |  |
| **Interpeak Latency** | | | |
| **Models** | **Deviance Information Criterion (DIC)** | | **Best Model** |
| **Null:** Latency ~ Intensity | 2105.32 | |  |
| **Additive:** Latency ~ Sociality + Sex + Intensity | 2102.80 | |  |
| **Full:** Latency ~ Sociality * Sex * Intensity | 2120.39 | |  |
| **Full + BM:** Latency ~ Sociality * Sex * Intensity + BM | 2121.59 | |  |
| **Amplitude Ratio** | | | |
| **Models** | **Deviance Information Criterion (DIC)** | | **Best Model** |
| **Null:** Amplitude ~ Intensity | 454.42 | |  |
| **Additive:** Amplitude ~ Sociality + Sex + Intensity | 450.10 | |  |
| **Full:** Amplitude ~ Sociality * Sex * Intensity | 412.94 | |  |
| **Full + BM:** Amplitude ~ Sociality * Sex * Intensity + BM | 413.89 | |  |
| **BIC Normalized Latency** | | | |
| **Models** | **Deviance Information Criterion (DIC)** | | **Best Model** |
| **Null:** Latency ~ ITD | 1402.96 | |  |
| **Additive:** Latency ~ Sociality + Sex + ITD | 1396.29 | |  |
| **Full:** Latency ~ Sociality * Sex * ITD | 1347.33 | |  |
| **Full + BM:** Latency ~ Sociality * Sex * ITD + BM | 1347.86 | |  |
| **BIC Relative Amplitude** | | | |
| **Models** | **Deviance Information Criterion (DIC)** | | **Best Model** |
| **Null:** Amplitude ~ ITD | 11859.88 | |  |
| **Additive:** Amplitude ~ Sociality + Sex + ITD | 11856.47 | |  |
| **Full:** Amplitude ~ Sociality * Sex * ITD | 11855.36 | |  |
| **Full + BM:** Amplitude ~ Sociality * Sex * ITD + BM | 11855.70 | |  |
